# Activity-Guided Proteomic Profiling of Proteasomes Uncovers a Variety of Active (And Inactive) Proteasome Species

**DOI:** 10.1101/2023.03.30.534963

**Authors:** Manisha Priyadarsini Sahoo, Tali Lavy, Indrajit Sahu, Oded Kleifeld

**Author notes:** Current address: Division of Medical Research, Faculty of Medical & Health Sciences, SRM University, Chennai 603203, India. These authors contributed equally to the manuscript (Manisha Priyadarsini Sahoo, Tali Lavy).

## Abstract

Proteasomes are multi-subunit, multi-catalytic protein complexes present in eukaryotic cells that degrade misfolded, damaged, or unstructured proteins. In this study, we used an activity-guided proteomic methodology based on a fluorogenic peptide substrate to characterize the composition of proteasome complexes in WT yeast, and the changes these complexes undergo upon the deletion of Pre9 (Δα3) or of Sem1 (ΔSem1).

A comparison of whole-cell proteomic analysis to activity-guided proteasome profiling indicates that the amounts of proteasomal proteins and proteasome interacting proteins in the assembled active proteasomes differ significantly from their total amounts in the cell as a whole. Using this activity-guided approach, we characterized the changes in the abundance of subunits of various active proteasome species in different strains, quantified the relative abundance of active proteasomes across these strains, and charted the overall distribution of different proteasome species within each strain. The distributions obtained by our mass spectrometry-based quantification were markedly higher for some proteasome species than those obtained by activity-based quantification alone, suggesting that the activity of some of these species is impaired. The impaired activity appeared mostly among 20S^Blm10^ proteasome species which account for 20% of the active proteasomes in WT.

To identify the factors behind this impaired activity, we mapped and quantified known proteasome-interacting proteins. Our results suggested that some of the reduced activity might be due to the association of the proteasome inhibitor Fub1. Additionally, we provide novel evidence for the presence of non-mature and therefore inactive proteasome protease subunits β2 and β5 in the fully assembled proteasomes.

**Significance Statement:** Proteasomes, essential protein complexes in eukaryotic cells, degrade misfolded, damaged, or unstructured proteins. Here we present an activity-guided proteomic method to characterize the composition and abundance of proteasomes. When applied to yeast proteasomes, this method revealed discrepancies between proteasome distributions determined by mass spectrometry and peptidase activity. This implies that a substantial portion of the proteasomes may exhibit reduced activity. Our findings indicate that these changes in proteasome activity could be linked to proteasome inhibition by Fub1. Furthermore, we identified signature peptides that indicate incomplete maturation of some of the β2 and β5 proteolytic subunits in fully assembled proteasomes, suggesting that proteasome core particle assembly can proceed even without the complete maturation of all β subunits.

## Introduction

The proteasome is a key player in the ubiquitin proteasome system of all eukaryotic organisms. It is responsible for the controlled degradation of proteins in almost all types of cells. The proteasome is an energy-dependent self-compartmentalized protease complex present both in the cytoplasm and nucleus of the cell. It degrades ubiquitin-modified, damaged, unfolded, and misfolded proteins in a well-regulated manner. In eukaryotic cells, the proteasome population predominantly contains three types of proteasomes – 30S, 26S and 20S (Figure 1A top). In addition to these main proteasome complexes, the 20S can form a complex with the proteasome activator Blm10 (1)-termed here 20S^Blm10^ (Figure 1A bottom).

**Figure 1.**
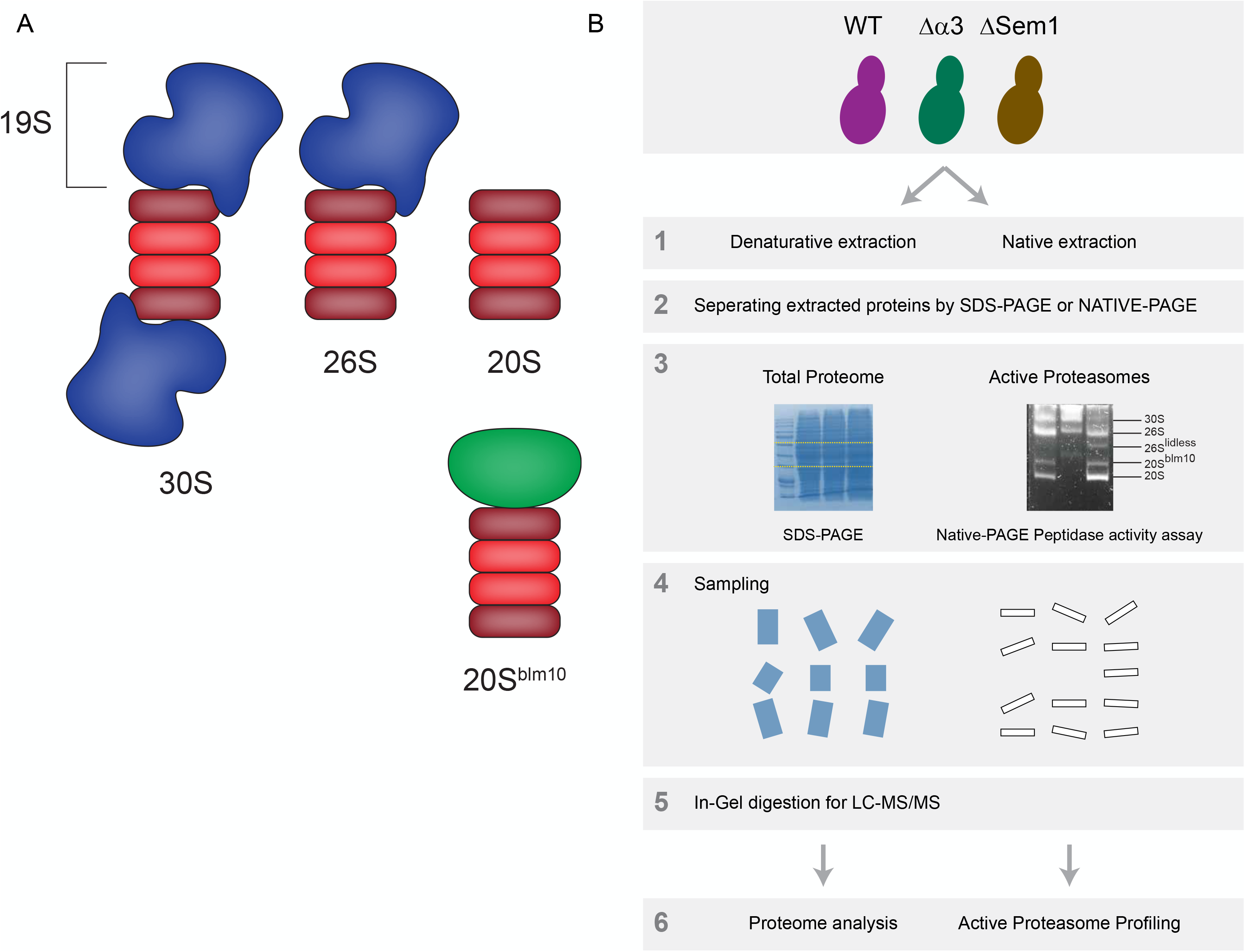
Activity-based proteomics to study active proteasome structures. **A**. The major types of proteasomes: 30S, 26S, 20S and 20S^Blm10^. The core particle 20S-CP is in red, the regulatory particle – 19S-RP is in blue, and Blm10 - the proteasome activator is in green **B**. Analysis workflow

Structurally, all these active forms of the proteasome contain a common core particle (CP) called 20S. The 30S proteasome contains two regulatory particles – 19S sub-complexes at both sides of its 20S core; the 26S proteasome contains one 19S at a single side of its 20S; and the 20S proteasome does not contain any regulatory particles. The 19S regulatory particle (RP) is mainly responsible for recognizing poly-ubiquitinated proteins, releasing ubiquitin chains, unfolding substrate proteins, and regulating the entry of the substrate into the 20S complex. The subunits of the 19S can be divided into subcomplexes termed ‘base’ and ‘lid’(2). The 20S core particle is the catalytic chamber that degrades the substrate into short peptides and amino acids with the help of the six catalytic subunits within it (3). The distribution of the different active proteasome species may vary under differing conditions (4). Attempts to determine this distribution using a variety of methods have led to different results. For example, SDS-polyacrylamide gel electrophoresis (PAGE) immunoblot analysis of the total amounts of proteasome subunits in *S. cerevisiae* suggested that the majority of proteasomes are present as 26S or 30S (5), while a proteomics study based on affinity-tagging and purification of proteasomes from mammalian cells showed that around 50% of all proteasome complexes are 20S (6).

Mass spectrometry (MS)-based structural and protein-interactions analyses of the proteasome holocomplex have been the focus of many research studies (7). Detailed structural characterization of the proteasome complex begins with its purification. Since the proteasome is an active protease complex, the challenge is to preserve its enzymatic activity for further functional studies whilst still obtaining a high purification yield. Given the high heterogeneity of proteasomes and the differences in stability and dynamics of proteasome interacting proteins (PIPs), the experimental conditions and the type of strategy used in the purification process highly influence the purity and composition of complexes obtained (30S, 26S, 20S proteasomes, and 20S with other activators), as well as the nature of the interactors purified (specific, stable, and dynamic).

Studies of native forms of yeast proteasome holocomplexes have been performed using 2D-gel electrophoresis by blue native PAGE followed by SDS-PAGE (8). This strategy, however, suffers from the reproducibility and complexity issues common to 2D-gel based methods. Alternatively, affinity purification methods such as immunoprecipitation, immunochromatography, and epitope tagging strategies, although they are costly and technically challenging, can generate a high yield of pure and functional proteasome complexes from a limited amount of starting material. Several affinity purification approaches have been developed over the past decade to purify proteasome complexes, and these approaches differ in several ways: the bait protein employed to catch the proteasome complexes (9), the use of an epitope tagged proteasome subunit (10), or the use of antibodies against proteasome subunits (11). In addition to the above one-step purification approaches, tandem purification strategies have also been developed to ensure the quality of the purified proteasome complexes for downstream studies (12).

When the isolated proteasome complexes by above methods are combined with MS-based proteomics, they can provide additional insight in relation to the dynamic variations of the different proteasome complexes and their PIPs networks (13). Prior to MS based analysis, the proteasome complexes can be chemically cross-linked before cell lysis, followed by one-step or tandem affinity purifications (10). For quantitative analysis of the different proteasome complexes and the PIP either chemical labeling (14), SILAC labeling (12), or label free (6) approaches were followed. Beyond the identification of the proteasome complex interactors, these proteomic studies enable the characterization of PTMs on proteasome complex subunits and possibly on their interacting partners. This quantitative proteomics could moreover allow the identification of transient and labile interactors, the measurement of proteasome complex dynamics in terms of composition and subcellular localization, and the determination of proteasome complex stoichiometry.

While the proteomic studies of proteasome complexes described above were successful, a lot of information may have been missed due to technical limitations. Many of these studies were based on affinity-based purification of the proteasomes, and as such their outcome was highly dependent on the specific proteasome subunit used for the purification (either by the addition of an affinity tag or by a specific antibody). Blue native PAGE is a simple method to separate and analyze protein complexes, however it cannot provide information on proteasome activity. To overcome this gap in existing information, we set out to develop a simple proteomic methodology that is based on common proteasome peptidase activity assays. We applied this methodology to study the changes in the yeast proteasome in two yeast strains generated by the deletion of two integral subunits of 26S proteasomes:Suppressor of Exocytosis Mutation 1(Sem1) and α3 (Pre9), hereafter referred to as Δsem1 and Δα3 respectively. Sem1 is a multifunctional and intrinsically disordered small (8 kDa) acidic protein in *Saccharomyces cerevisiae* that is conserved among eukaryotes. It is one of the 19S lid proteins (15) and is also crucial in its assembly (16). Sem1, while not an essential gene, regulates exocytosis and pseudohyphal differentiation in yeast (17). In *Saccharomyces cerevisiae*, the α3 subunit gene is the only non-essential CP gene (18, 19). Yeast lacking α3 form an alternative proteasome by substituting an additional α4 subunit where α3 normally resides (19). Interestingly, similar alternative proteasomes that contain 2 copies of α4 without any α3 have also been observed in mammalian cells under certain stress conditions (20).

To examine changes in the proteasome among these strains relative to the wild-type (WT), we performed side-by-side proteomic characterizations of the whole cell and activity-guided proteasome profiling (Figure 1B). Our analyses indicate that the amount of active proteasome complexes in the cells does not necessarily correspond to the cellular abundance of proteasome subunits. We determined the relative amount of each active proteasome specie within each of the studied strains, investigated the distribution of various known proteasome-interacting proteins (PIPs), and assessed the maturation of the proteolytic subunits of the proteasome across different active proteasome complexes.

## Results

### Workflow

In-gel activity assay of the proteasome is commonly used for characterization of the levels of different active proteasomes in the studied sample. It is based on a clear native gel that is immersed in a buffer containing a fluorogenic peptide substrate. The peptidase activities are visualized in the gel under a UV illuminator (21). We take advantage of this assay’s simple and quick proteasome separation and excise the active proteasome bands for in-gel digestion followed by LC-MS/MS analysis (Figure 1B).

The isolation of active proteasome species from a native gel has several advantages: it allows direct characterization of the different proteasomal species obtained under native conditions without the need to isolate them by other methods which may lead to biased results, such as purification of over-expressed and/or tagged proteasomal species; it allows concurrent characterization of different proteasomal species present in the same sample; the usage of quantitative proteomics enables an accurate determination of the relative abundance of the proteasomal species which does not depend on the efficiency of the activity assay, hence overcoming biases resulting from different efficiencies of the activity assay for different proteasomal species (21).

### Whole proteome analysis shows elevation in proteasomal proteins in mutant strains relative to the WT

As a reference for our activity-based proteasome characterization, we utilized label-free proteomics to compare the whole proteomes of WT, ΔSem1, and Δα3 strains, and to determine the expression level of the different proteasome subunits.

Following common MS/MS data analysis steps, a total of 2597 proteins were identified and quantified (Supplement Table S1). PCA of LC-MS/MS data of 3 repeats from each strain shows distinct clusters for each strain, indicating that the strains differ with respect to the whole proteome content (Supplement Figure S1).

Following multi-sample ANOVA statistical analysis, 1228 proteins with significant abundance changes were identified. Hierarchical clustering of these 1228 proteins shows that they can be grouped into 6 distinct clusters with different expression profiles across the three strains (Figure 2A). Go Terms enrichment of these proteins (Supplement Table 2) reveals several defined categories with significant changes. As shown in Figure 2A, one of these categories is proteasomal proteins, whose levels are elevated in the mutants with respect to the WT. We performed a two-way T-test to further compare each of the mutant strains with the WT (Figure 2B and 2C). Each of the mutant strains showed significant differences in the expression of around 600 proteins relative to WT. The amount of most proteasomal proteins is elevated in both mutants when compared against the WT (Figure 2B and 2C top panels). In order to test the overall changes in the 19S and 20S subcomplexes, we summed the LFQ intensities of the proteins that make up these subcomplexes in each of the mutants and compared them to the WT values (Figure 2B and 2C bottom panels). This indicates that the increase in the expression level of the proteins that make up 19S in the Δα3 strain is higher relative to the expression of the proteins that make up the 20S (Figure 2B bottom). In the case of ΔSem1, the change (relative to the WT) in the amount of the proteins that make up the 19S or the 20S is about the same (Figure 2C bottom).

**Figure 2.**
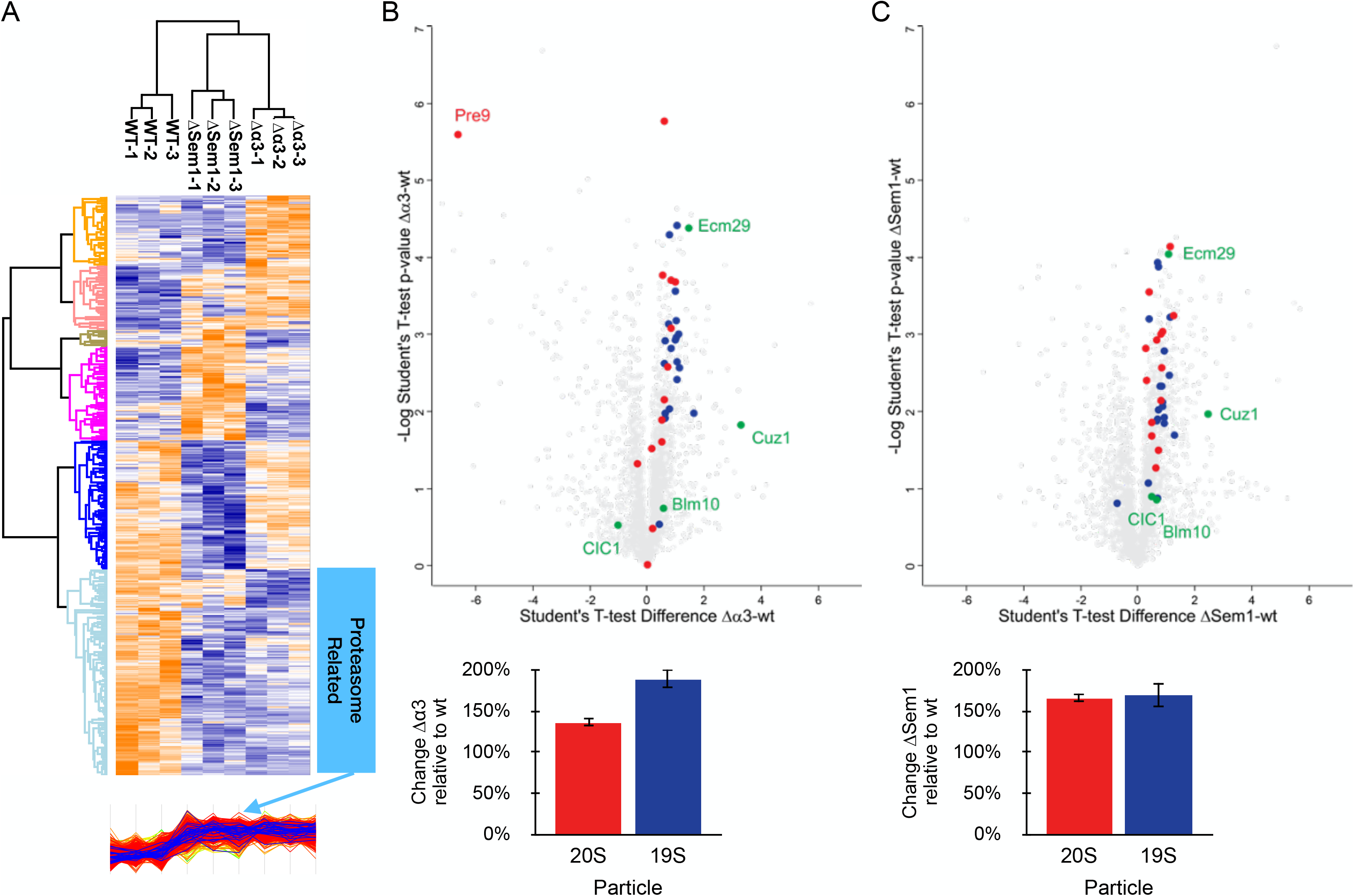
Whole cell proteomic data analysis of WT, ΔSem1 and Δα3. **A.** Hierarchical clustering of the 1228 proteins that show significant abundance changes across the different strains. Lower cluster (marked) in light blue is of proteolysis-related proteins. **B.** Top panel: T-Test results of the comparison of WT and Δα3 strain. CP subunits are shown in red and RP in blue. Known PIPs are shown in green. Bottom panel: average fold change of proteasome CP and RP subunits in Δα3 relative to WT. **C.** Top panel: T-Test results of the comparison of WT and ΔSem1 strain. CP subunits are shown in red and RP in blue. Known PIPs are shown in green. Bottom panel: average fold change of proteasome CP and RP subunits in ΔSem1 relative to WT.

### Proteomic characterization of the active-proteasome complexes

The abundance of the different proteasome subunits in the whole cell lysate does not necessarily represent their levels within the different active proteasome complexes. To characterize the proteomic changes in the active proteasome complexes in these strains we used native gel, excised the active bands of each strain, and submitted them to LC-MS analysis following in-gel digestion (as in Figure 3A). If no active band was observed we used the WT strain bands (analyzed on the same gel) as a reference and excised the corresponding same regions from all lanes (Supplement Table S3). Next, to characterize strain-related proteasomal content, we used LFQ and compared the abundance of all active proteasomes of each mutant relative to those of the WT (Figure 3A). In the case of Δα3 strain, the relative abundance of all 20S subunits except Pre9 (α3 itself) and Pre6 (α4) is significantly lower compared to the WT (Figure 3B). An additional copy of Pre6 (α4) compensates for the absence of α3 in this strain and forms a complete α-ring (19, 22). Therefore, its unique abundance change among the other 20S subunits is anticipated. In the ΔSem1 strain, the relative abundance of the 19S subunits is lower on average compared to the WT, while the abundance of the 20S subunits is higher (Figure 3C).

**Figure 3.**
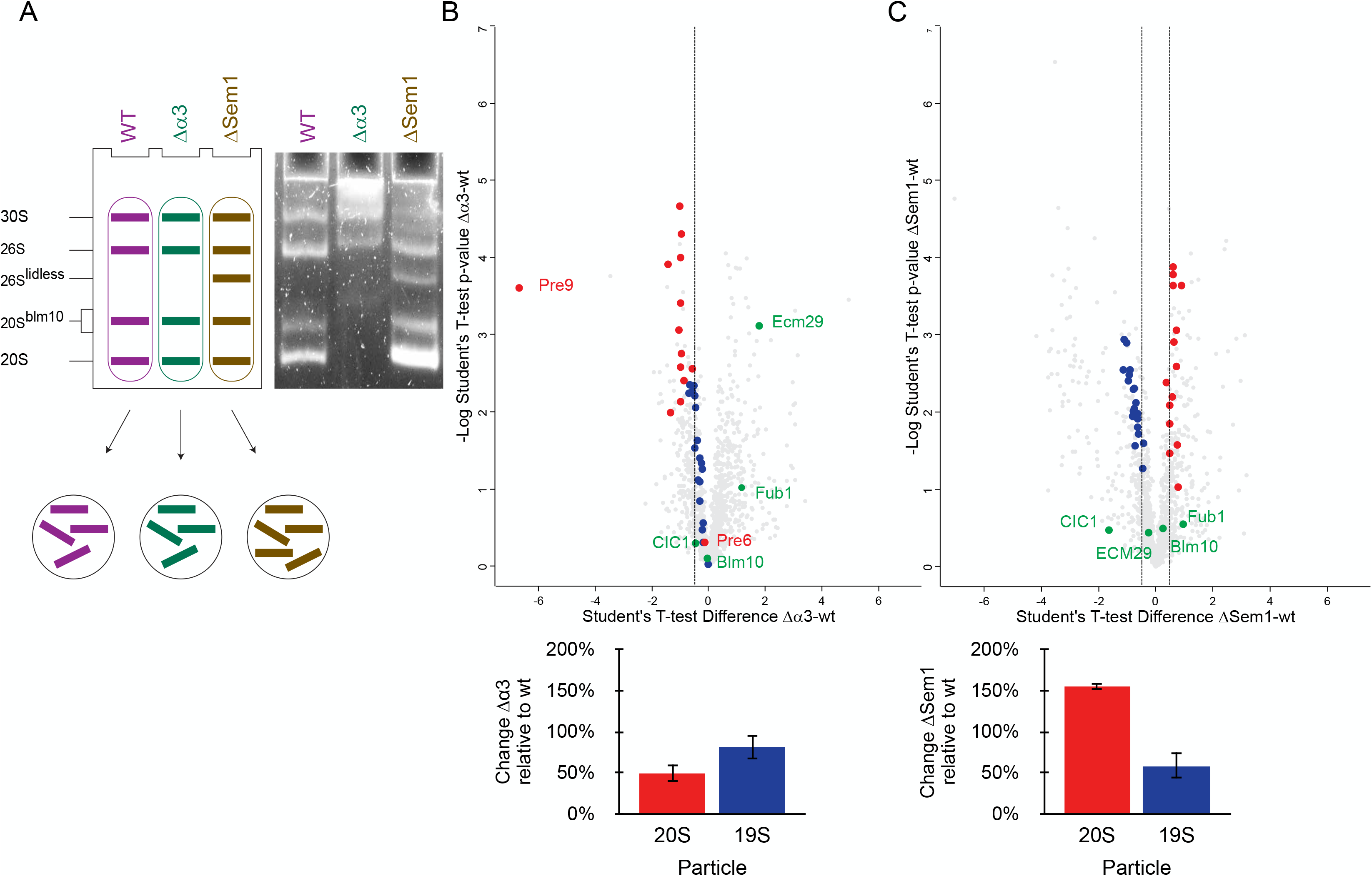
Activity-guided proteomic profiling of total active proteasome content. **A.** Activity-guided analysis strategy for relative comparison of the total amount of proteasome complexes between strains. **B.** Changes in proteasome subunits content of different proteasome complexes in Δα3 relative to WT. Top panel: T-Test results of the comparison of WT and Δα3 strain. CP subunits are shown in red and RP in blue. Known PIPs are shown in green. Bottom panel: average fold change of proteasome CP and RP subunits in Δα3 relative to WT. **C.** Changes in proteasome subunits content of different proteasome complexes in ΔSem1 relative to WT. Top panel: T-Test results of the comparison of WT and ΔSem1 strain. CP subunits are shown in red and RP in blue. Known PIPs are shown in green. Bottom panel: average fold change of proteasome CP and RP subunits in ΔSem1 relative to WT.

This is expected, given that Sem1 is a stoichiometric component of the 19S and hence is essential for efficient lid assembly (23). This correlates with the in-gel peptidase activity assay, which shows a decrease in the amount of active 26S and 30S in ΔSem1 relative to WT (Figure 3A and Figure S2). Sem1 is notably absent from these graphs, since it was identified by a single unique peptide due to its small size and typical tryptic digestion pattern, leading to its label-free quantification (LFQ) not being calculated. This single and unique Sem1 peptide was identified only in Δα3 and WT (in all 3 biological repeats).

As for known PIPs, the levels of Blm10 (24) and Ecm29 (25) in ΔSem1 were similar to the WT (Figure 3C). This is in contrast to the relative abundance changes of Ecm29 in the whole cell proteomic comparison (Figure 2C), which shows that it is significantly enriched in ΔSem1 compared to WT despite the Blm10 abundance not changing significantly (Figure 2C). Our results indicate that Ecm29 levels are significantly higher in Δα3 proteasomes compared to WT while Blm10 levels are similar (Figure 3B). In this case the levels in the proteasome complexes correlate well with the whole cell levels that show similar trends (Figure 2B). Ecm29 tethers the 20S to the 19S and preserves their interaction when ATP is absent (25, 26), is known to recognize faulty proteasomes (27, 28), and plays a role in proteasome changes under oxidative stress (29). In Δα3 there is a problem with 20S assembly which leads to a high ratio of fully assembled 19S to 20S. This can be observed by in-gel peptidase activity results showing that in Δα3 the proportion of 30S and 26S proteasome complexes compared to 20S is higher than in WT cells (Supplement Figure S2). To ensure that Δα3 strain retains the appropriate levels of functional 30S and 26S proteasomes, it is likely that Ecm29 has been upregulated compared to WT and is now associated with these proteasome complexes. Other PIPs like the adaptor protein Cic1(30) and the proteasome inhibitor Fub1 (31, 32) were identified and quantified but did not show significant changes between WT and Δα3 or ΔSem1. It is worth noting that Fub1 was only identified and quantified in the activity-guided proteomic profiling of the proteasomes (Figure 3B and 3C) and was not quantified in the whole cell proteome analysis. In contrast, Cuz1, the Cdc48-associated UBL/Zn-finger protein, known to deliver ubiquitinated substrates to the proteasome (33, 34), showed higher expression in Δα3 and ΔSem1 relative to WT in the whole proteome analysis (Figure 2B and 2C, respectively). However, its identification by the activity-guided proteasome profiling was much weaker and did not allow for quantification under the analysis settings that were used (Supplement table S3). Taken together, these results indicate that the expression levels of proteasomal proteins and PIPs in cells differ significantly from their amounts in the assembled active proteasomes.

### Relative amount of active proteasome species in each strain

We performed a pairwise LFQ comparison of the different active proteasome complexes relative to the 26S species (Figure 4A and Supplement Table S4) to characterize the changes in proteasome subunits and PIPs in each active proteasome species within a strain. The comparison of WT 26S to 30S proteasomes showed that the amount of CP subunits is higher in the 26S species than in the 30S species (Figure 4B), with an average fold change of ∼2. The intensities of the RP subunits were very similar in both species, and since the 30S species contains two RP sub-complexes, the results suggest that there is twice as much 26S than 30S in WT cells. In the comparisons of WT 26S to 20S (Figure 4C) and 26S to 20S^Blm10^ (Figure 4D), the expected absence of the 19S subunits from the 20S samples is reflected. In the 20S sample there are more 20S subunits (1.33 fold) than in the 26S sample, while in the 20S^Blm10^ sample there are less (0.77 fold). Abundance changes were also observed for Ecm29, but its lower level in the 26S relative to the 30S is anticipated given its roles in the assembly of the 20S and 19S species and in proteasome quality control. The higher amount of Blm10 in the 26S sample suggests that some of the 26S proteasomes are hybrid proteasomes that have Blm10 on one side of the 20S and 19S on the other as reported before(1). This indicates that these two species (26S, and 26S^Blm10^) could not be resolved in our activity gel.

**Figure 4.**
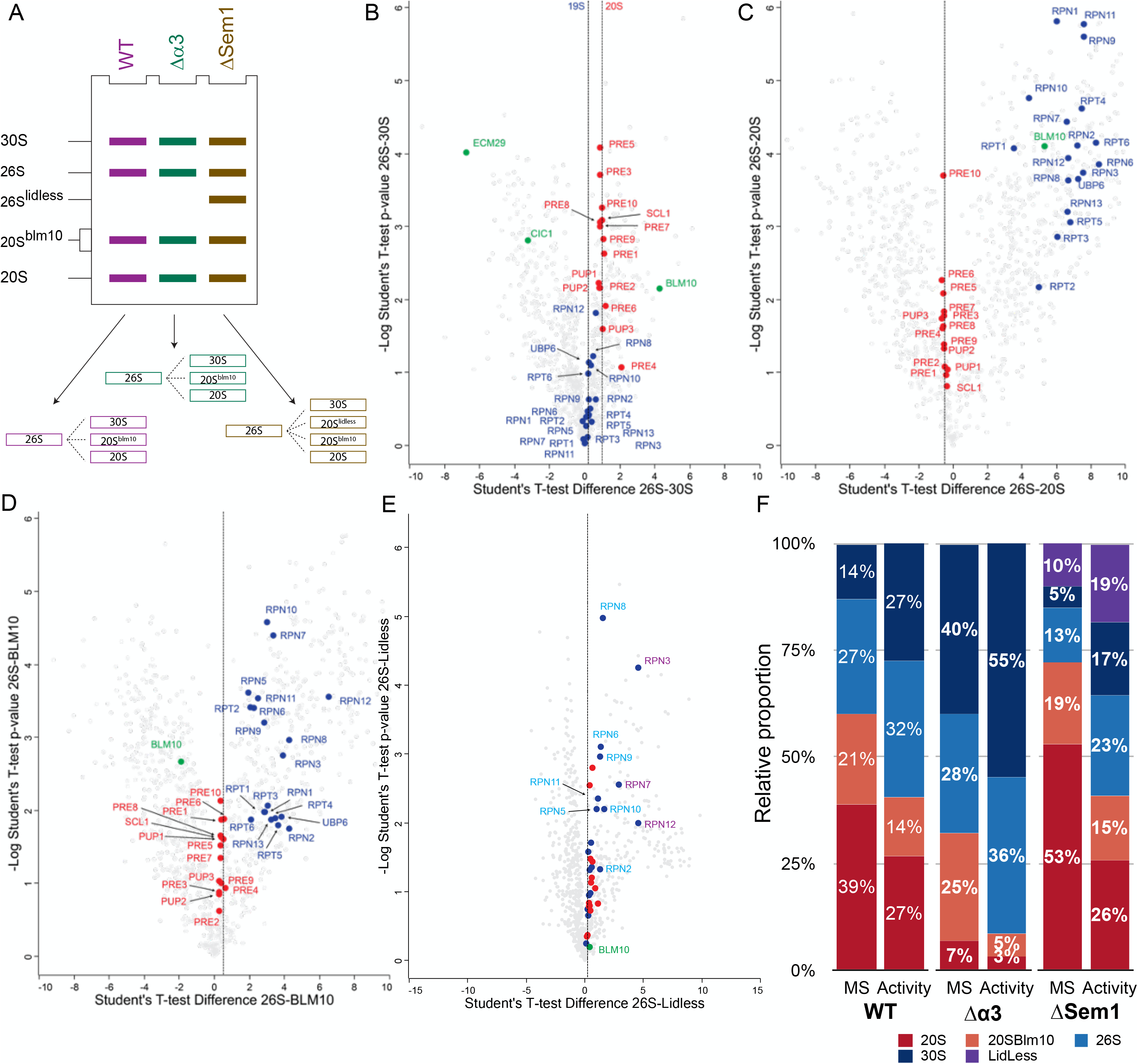
Distribution of proteasome complexes within each strain. **A**. Analysis strategy of proteasome complexes within each strain based on the relative comparison to the 26S band of each strain. **B**. Comparison of proteins identified in WT 26S band to those that were identified in 30S. CP subunits are shown in red and RP in blue. Known PIPs are shown in green. **C**. Comparison of proteins identified in WT 26S band to those that were identified in 20S. CP subunits are shown in red and RP in blue. Known PIPs are shown in green. **D**. Comparison of proteins identified in WT 26S band to those that were identified in 20S+Blm10 band. CP subunits are shown in red and RP in blue. Known PIPs are shown in green. **E.** Comparison of proteins identified in ΔSem1 26S band to those that were identified in ΔSem1 26S-Lidless band. CP subunits are shown in red and RP in blue. Rpn subunits that show 2-3 fold change are labeled cyan and those that show >4 fold change are marked in purple. Known PIPs are shown in green. **F**. The relative distribution of active proteasome complexes in the different yeast strains. For each strain the relative distribution of proteasome content calculated from LFQ analysis of active proteasome bands is shown on the left (marked as “MS”) and the distribution calculated based on active proteasomes band intensity in native gels is shown on the right (marked “Activity”).

Similar comparisons were also conducted for the active proteasomes of Δα3 (Supplement Figure S3 and Supplement Table S5) and ΔSem1 (Supplement Figure S4 and Supplement Table S6). In the case of the ΔSem1 strain, we also included a comparison of the lidless 26S proteasome, which is unique to this strain (Supplement Figure 5A). This particular proteasome species displayed distinct ratios for various proteasomal subcomplexes and subunits, as shown in Figure 4E and Supplement Figure S5B. Compared to the lidless 26S, the 20S subunits and the 19S base Rpt1-6, Rpn1, Rpn13, and Ubp6 were approximately 1.4 times more abundant in the 26S. As expected, most of the lid subunits Rpn5, Rpn6, Rpn8, and Rpn9 showed significantly higher abundance in the 26S relative to the lidless 26S (about 2-3 times higher). The same trend of higher relative abundance in the 26S was observed for the base subunits Rpn2 and Rpn10, possibly reflecting Rpn2’s position and its contacts with the lid subunits (35) and Rpn10’s role as the hinge between the base and lid (35, 36). The most significant changes in abundance were found for Rpn7, Rpn12, and Rpn3, which were all significantly higher in the 26S band. These results match Sem1’s position in the proteasome structure (Supplement Figure 5C) and supports its role as a chaperone for lid assembly, which involves the binding of Rpn7 and Rpn13 before they associate with the rest of the lid followed by the binding of Rpn12 to complete the lid assembly(23, 37).

In the next step, we conducted proteomic analyses of the different active proteasome complexes to determine the relative amount of each type of proteasome within each strain. For this purpose, we used the LFQ intensities of all the 20S subunits as a reference since they are common to all active proteasome complexes. We calculated the relative proportion of each active proteasome species out of the total active proteasomes in each strain (Figure 4F). The analysis revealed that in the WT strain, the majority of proteasomes in the cell are 20S (Figure 4F left). Its share of the total active proteasomes (39%) is equivalent to the amounts of the 26S and 30S proteasomes combined. Furthermore, the relative amount of the WT 26S proteasomes (27%) is approximately twice that of the WT 30S proteasomes (14%). Interestingly, WT 20S^Blm10^ comprises about one-fifth of the total proteasomes in the cell and its amount is higher than that of the WT 30S proteasomes. The MS-based distribution of active proteasomes differs significantly from the proportions based on the in-gel peptidase activity of the WT sample (Figure 4F and supplement Table S7). According to the activity assay, most of the WT proteasomes are 26S (32%), while the 30S and 20S account for 27% of the total each, and the smallest portion consists of 20S^Blm10^, which makes up only 14% of the total.

The MS-based quantification of Δα3 proteasomes (Figure 4F middle) reveal a very high proportion of 30S (40%) and a very small portion of 20S proteasomes (7%). The rest of active proteasomes are split almost equally between 26S (28%) and 20S^Blm10^ (25%) proteasomes. By peptidase-activity-based quantification the relative share of 30S and 26S is much larger and reaches over 90% of the proteasomes while 20S^Blm10^ portion is only 5% and the 20S is only 3%. These quantifications highlight again the 20S assembly issues due to the lack of α3. The reduction in fully assembled 20S in this strain may lead to a large excess of assembled 19S that readily reacts with any available 20S, thus most of the active Δα3 proteasomes contain RP.

For ΔSem1, the MS-based quantification (Figure 4F right) indicates that more than half of the proteasomes in this strain are 20S (53%) and almost none are 30S proteasomes (5%). The remainder is split between 20^Blm10^ (19%), 26S (13%) and lidless-26S (10%). This confirms Sem1’s role in the lid and RP assembly, which causes a shortage in assembled 19S and affects the association of the CP and RP. By peptidase-activity-based quantification the relative share of 30S and 26S is much larger and reaches over 50% of the proteasomes while the 20S and 20S^Blm10^ portions are reduced to 26% and 15% respectively.

### Relative amount of PIPs in the different proteasome complexes

Reduction and enhancement of proteasomal activity can be caused by the presence of additional proteins that can bind specifically to the proteasome and alter its activity (1, 27, 31, 38). Therefore, we used the same type of quantitative analysis described above (Figure 4F) to monitor the distribution of the known PIPs. In our dataset, among a multitude of ubiquitin-proteasome system-related proteins, we identified and quantified 19 PIPs known to play essential roles in proteasome assembly, storage, or activity modulation. (Supplement Table S8). Of these, we focused on PIPs that are known to affect proteasome activity (Figure 5) and might explain the results described above (Figure 4F). Based on all the LFQ intensities that were obtained for particular PIP in each proteasome complex we calculated its distribution across those complexes as described in the methods section.

**Figure 5.**
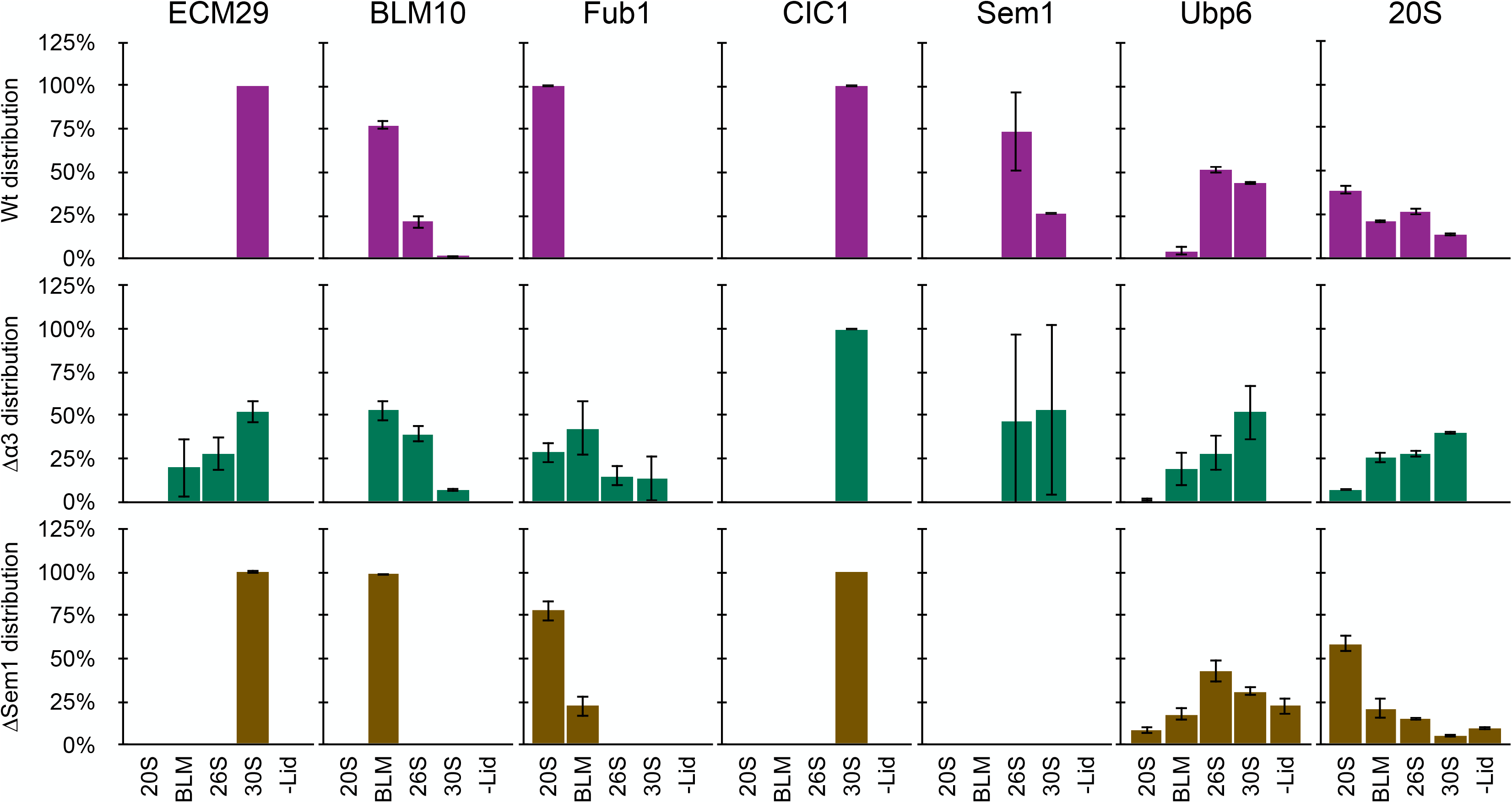
Distribution of known proteasome-interacting-proteins. The relative amount of known PIPs in each of the active proteasome complexes was calculated from the LFQ analysis of active proteasome bands. The distribution of complexes is indicated by the 20S (right). The distribution of the PIPs among the various proteasome complexes is shown in purple for WT, in green for Δα3 and in gold for ΔSem1. The 20S and Ubp6 distributions are shown as references for the distribution of the active proteasomes and RP, respectively.

Our analysis revealed that only one protein, Cic1, was exclusively associated with the 30S proteasome in all three strains. The specific binding of Cic1 to the 30S proteasome was previously demonstrated using a tagged version of Cic1 (30). However, this binding was not linked to Cic1’s suggested function as a proteasomal adaptor for specific types of substrates, nor to its localization within the nucleus (30). Ecm29 specifically interacts with the 30S proteasome in both WT and ΔSem1. This interaction corresponds to the reported binding of Ecm29 to both the 20S and 19S proteasomes that link them together (25). Yet, previous research using 2D native-PAGE showed that Ecm29 can also interact with the 26S proteasome (26). However, this discrepancy may be due to the different methodologies used, and also since the reported interaction of Ecm29 with 26S proteasomes was shown for purified proteasomes. Despite the relatively low abundance of 30S proteasomes in ΔSem1 (Figure 4F), Ecm29 still specifically interacted with these proteasomes. This may be because the lack of Sem1 generates incomplete proteasomes (such as Lidless 26S) that cannot properly bind Ecm29, despite the higher abundance of Ecm29 in ΔSem1 cells (Figure 2C). Surprisingly, Ecm29 was detected in all proteasome species in Δα3, except for the 20S proteasome. This could be explained by the unique nature of Δα3 and the reported ability of Ecm29 to function as a quality control factor by binding to aberrant proteasomes (such as Δα3) and ensuring their inhibition by gate closure (27).

Blm10 is known to bind to the proteasome CP in both 20S and 26S (39). In all strains tested, Blm10 was mostly or exclusively detected in the 20S^Blm10^ proteasome. Unlike a previous report suggesting that the majority of wild-type (WT) Blm10 is bound to the 26S proteasome (1), our findings indicate that only 25% of WT Blm10 is bound to 26S, while most Blm10 is in 20S^Blm10^. Notably, in the ΔSem1 strain, Blm10 was only detected in 20S^Blm10^, likely due to the assembly defect of the RP that reduces the 26S proportion compared to WT. In contrast, in the Δα3 strain, the proportion of Blm10 bound to 26S was higher than in WT, possibly due to the altered 20S assembly that leads to much less 20S in this strain than in WT.

Although Blm10 is typically described as a proteasome activator (1, 39), it has been reported to bind to both sides of the 20S and generate an inactive form of proteasome called Blm10_2_-CP (39). The formation of this Blm10_2_-CP was found to be enhanced in open-gate proteasome mutants such as α3 or α7 that lack their N-terminal domain (39). In Δα3, we observed a significant difference between the proportion of 20S^Blm10^ obtained based on chymotrypsin peptidase activity (5%) and that was determined by MS (25%) (Figure 4F). Therefore, it is possible that the gel region we assigned and excised as Δα3 20S^Blm10^ may have contained a mix of 20S^Blm10^ and the Blm10_2_-CP.

The proteasome inhibitor Fub1 has been less studied compared to other PIPs. Fub1 has been shown to physically interact with the CP subunits(32). When comparing the relative quantities of proteasomes and Fub1 in different strains (as depicted in Figure S6A and S6B, respectively) it is evident that Fub1 is more abundant in Δα3 strain, corroborating recent findings (31). In WT, Fub1 is exclusively associated with the 20S proteasome. However, in Δα3, Fub1 is present in both the standalone CP and CP-RP complexes, with a preference for 20S and 20S^Blm10^. This is consistent with previous findings for this strain(31). In the ΔSem1 strain, Fub1 is exclusively associated with 20S and 20S^Blm10^, in proportions that mirror the overall distribution of these proteasomes in this strain. Hence, the presence of Fub1 could also account for some of the observed divergence between the mass spectrometry-based and activity-based quantifications detailed above.

### Evaluation of proteasome proteolytic subunits activation

Most proteases, including the proteasome proteolytic subunits (β1, β2, and β5), are initially translated as inactive zymogens and must undergo proteolytic processing to become active. This self-catalytic process occurs during the final steps of CP assembly, after the two halves of the CP have been dimerized, and results in the exposure of the active site threonine of the proteolytic subunits (40). We wanted to test if the underestimation of some proteasome species based on their activity (Figure 4F) might be explained by certain alterations in the maturation process of β1, β2, and β5. However, typical bottom-up proteomics approaches, which rely on conservative peptide searches that assume fully specific tryptic digestion, are not able to detect endogenous proteolytic processing, such as the maturation of beta subunits. To identify proteasome proteolytic subunit activation events, we analyzed all active proteasome samples using peptide search that is based on semi-tryptic cleavage specificity (Supplementary Table S9). Through this method, we identified several signature peptides that span the cleaved propeptide domain (Figure 6A), providing insight into the activation status of the proteases in each complex. This type of analysis only looks at a small and specific group of peptides, so it’s likely that no signature peptide will be identified if the level of the studied complex is low. The MS/MS identifications of proteolytic beta subunits signature peptides had high scores and high peptide sequence coverage therefore provided conclusive evidence for their presence in the different samples (Figure S7 and S8). We also identified semi-tryptic signature peptides that indicate the maturation of β6 and β7 subunits.

**Figure 6.**
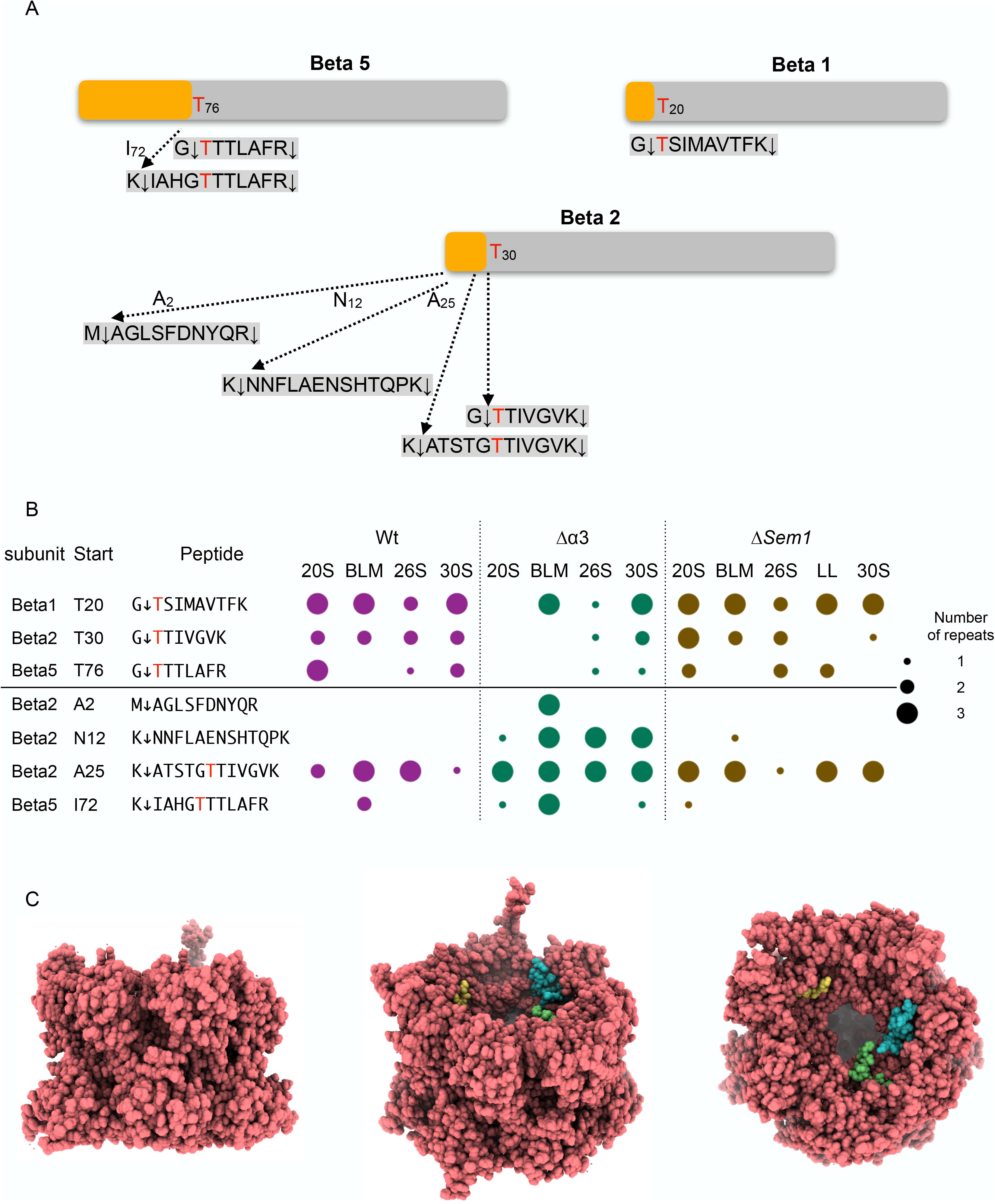
Activation of proteasome proteolytic subunits. **A**. Tryptic and semi-tryptic peptides that span the propeptide domain of the proteasome proteolytic subunits. The identification of these signatures indicates the activation status of the proteolytic subunits: β1, β2 and β5. The propeptide domain is shown in yellow and the active protease chain is shown in gray. The catalytic threonine is shown in red. **B**. The number of identifications of the different proteasome protease prodomain peptides among the various proteasome complexes. WT in purple, Δα3 in green, and ΔSem1 in gold. The size of the dot represents the number of biological repeats in which the peptide was identified. Peptides that indicate activation of the beta subunits are in lines above the partition line and peptides that indicate the presence of the uncleaved prodomain are below the line. **C**. Yeast proteasome structures that include proteolytic subunits propeptide domains. The structures are from (47) that were aligned and overlaid using ChimeraX(58). For clarity, only the 20S core (in red) is shown from different viewpoints. The remnants of prodomain β1 (residues Gly3 to Gly19, in cyan) and β2 (residues Asn18 to Gly29, in green) are from structure 5fgi and the remnant of prodomain β5 (residues Ile71 to Gly75, in yellow) from structure 5cz7.

As expected, active β1, β2, and β5 subunits were identified in all WT samples, except for β5 in 20S^Blm10^ samples (Figure 6B, top lines). A similar trend was observed for ΔSem1, in which we identified at least 2 different active proteolytic subunits in all proteasome complexes. We found less active proteases in the Δα3 complexes. This was most noticeable in the Δα3 20S sample, where there was no indication of any active β subunit. On the other hand, the signature peptides that indicate the presence of non-mature and inactive β subunit (Figure 6B bottom lines) were identified in all proteasome complexes and strains. Non-mature β2 (lacking the trypsin-like activity) was identified in all WT, ΔSem1 and Δα3 while non-mature β5 (lacking the chymotryptic-like activity) was identified in WT 20^Blm10^, ΔSem1 20S, and in all Δα3 complexes aside from the 26S. Of note, the presence of non-matured β2 and β5 was particularly prevalent in Δα3 20^Blm10^ where all 4 possible non-mature signature peptides appeared in all the biological repeats. In the CP assembly process, the autocatalytic cleavage of the β subunits prodomain is accompanied by the degradation of the proteasome-assembly factor UMP1(40, 41). We were able to identify UMP1 peptides only in some of the Δα3 proteasome samples (Supplement Table S9).

This fits well with the very low activity observed for this species (Supplement Figure S2) and its relative high abundance determined by MS (>25%, Figure 4F). Evidence for non-matured 20^Blm10^ was found for all 3 strains indicating that this may be a common character of this complex and may also contribute to its altered activity. Interestingly, structural modeling indicates the presence of the prodomains of the beta subunits does not alter the overall structure of the 20S proteasome. Furthermore, these additional chains can be easily accommodated within the 20S proteasome’s cavity. (Figure 6C).

## Discussion

The proteasome is the primary contributor to the proteolysis of most cellular proteins in eukaryotes. Consequently, any fluctuations in the levels of cellular proteasome complexes can disrupt protein homeostasis. To investigate such changes in active proteasome ratios within cells, we present a straightforward yet potent method for activity-guided, MS-based proteomic profiling of proteasome complexes. This method relies on peptide substrates, preserves the native and active forms of proteasome, and separates these various active complexes using clear native PAGE.

This concept is not new; in fact, it was one of the first methods used for the initial characterization of the proteasome as an ATP-dependent proteolytic complex (42). Over the years, the PAGE separation process and the activity measurements have been streamlined and standardized(43), so it is easy to implement this method in any biochemical or cell biology laboratory. Overall, this approach is simple, cost-effective, quick, and can be broadly applied to cultured cells or tissue samples as sources of proteasomes.

In this work, we used this method to determine the composition and relative abundance of each type of active proteasome in different yeast strains. In the WT strain, we found that the 20S proteasome was the most abundant, with a relatively large quantity of 20S^Blm10^ species also present. This relatively higher abundance of yeast 20S compared to 26S and 30S in the WT is in accordance with MS-based studies of the abundance of constitutive proteasomes in mammalian cells (6). Aberrations in CP or RP assembly, caused by Δα3 and ΔSem1 respectively, strongly affected the distribution of the different proteasomes.

Interestingly, in all of the studied strains we observed a discrepancy between the proteasome distribution determined by MS as opposed to that determined by monitoring in-gel peptidase activity (Figure 4F). This discrepancy was mostly noticed regarding the relative quantification of the 20S and 20S^Blm10^ species. The discrepancy between MS-based and activity-based quantifications is partly related to the limitations of the peptidase activity assay. While activity-based quantifications are useful in detecting active proteasomes, they do not necessarily provide accurate data on their relative amounts. The substrate access to the CP catalytic cavity is controlled by the α subunits ring, which surrounds the entry channel (44). The RP can open the entry channel, allowing the fluorogenic substrate to enter more easily through the CP gate in 26S and 30S proteasomes. In contrast, 20S and 20S^Blm10^ proteasomes typically require the addition of small amounts of detergent, like SDS, to promote gate opening (43). This is likely leading to altered quantification and overestimation of the 30S and 26S complexes. Furthermore, the in-gel peptidase activity assay is qualitative by nature and is read at a single time point, usually after a relatively long incubation time, which can lead to saturation and therefore inaccurate quantification.

To investigate other factors that might contribute to the observed reduction in proteasome activity of the 20S and 20S^Blm10^, we characterized and quantified the PIPs associated with each proteasomal species. This analysis provided clear and novel information regarding the distribution of known PIPs amongst the different active proteasome complexes. Our PIP mapping suggest that changes in proteasome activity, particularly of 20S and 20S^Blm10^, might be related to proteasome inhibition by Fub1. Recent research has revealed the mechanism by which Fub1 inhibits proteasomes in Δα3 (31), but one unanswered question is how it enters the CP cavity. Δα3 proteasome is known to have an open CP gate and therefore it was suggested that Fub1 recognizes proteasomes with an aberrant CP gate and enters through them into the matured CP, while simultaneously blocking all proteolytic subunits(31). Unlike Δα3, WT and ΔSem1 proteasomes have a typical CP entry gate which is not continuously open, which emphasizes the enigma regarding Fub1’s distribution. One possible answer, as suggested by Rawson et al (31), is that Fub1 might be part of a quality control mechanism that inhibits a subset of CP with abnormal entry gates. Another option is an alternative entry mechanism for CP with a closed entry gate.

Additionally, we identified signature peptides for the maturation of the proteolytic subunits of the proteasome, which surprisingly revealed the presence of non-cleaved and therefore inactive β2 and β5 subunits in the assembled proteasomes. These uncleaved subunits may also account for the reduced activity observed in the proteasomes. The autocatalytic cleavage of the proteolytic proteasomal β subunits is a critical event in the assembly of the CP (40, 45–47). This process takes place in the final step of assembly and requires the dimerization of two half-proteasome particles, each composed of a full α ring and a full β ring (40). This sequence of events ensures that the CP chamber is sealed off from the cellular environment before its proteolytic activity is initiated.

The presence of non-mature signature peptides indicate the presence of full or partial β2 and β5 propeptide domains. It has been shown before that yeast strains carrying a single mutation at the catalytic threonine of β1 or β2 or both are still viable, and that their proteasomes can accommodate the remnants of the propeptide domain(47, 48). However, outright deletion of the β5 propeptide domain or disruption of its cleavage process leads to significant phenotypic variations(47). Here we show, for the first time, that proteasomes with disrupted or incomplete processing of β2 and β5 propeptide domains are present in WT yeast. Additional studies are required to determine the mechanism that allows proteasome assembly despite the incomplete propeptide removal.

In conclusion, we have demonstrated that our activity-guided proteomic method for studying the proteasome is a valuable addition to the proteasome research toolbox. It provides new and detailed information that can be obtained by the current, commonly-used proteasome characterization methods.

## Materials and Methods

### Reagents

All reagents were ordered from Sigma unless specified otherwise.

### Yeast strains and growth conditions

All single yeast strains were purchased from Euroscarf:

WT: MATa BY4741; MATa; ura3Δ0; leu2Δ0; his3Δ1; met15Δ0 (Euroscarf # <u>Y00000</u>)

Δα3: MATa ura3Δ0; leu2Δ0; his3Δ1; met15Δ0, YGR135w::kanMX4 (Euroscarf # <u>Y04765</u>)

ΔSem1: MATa; ura3Δ0; leu2Δ0; his3Δ1; met15Δ0; YDR363w-a::kanMX4 (Euroscarf # <u>Y04200</u>)

Yeast strains were cultured at 30 °C in yeast extract peptone dextrose (YPD) or synthetic media (SD) according to the requirements of different experiments. YPD medium consisted of 1% yeast extract, 2% Bacto Peptone (Difco) and 2% dextrose(49). Synthetic media consisted of 0.7% Yeast Nitrogen Base supplemented with amino acids, adenine, and 2% dextrose(50).

### Sample preparation for whole proteome analysis

Yeast cells sample preparation for whole proteome analysis was done as described before (51, 52). In brief, 5-mL yeast cultures of each strain were grown until OD_600_ reached 1.5. The cells were collected by centrifugation, washed twice with cold double-distilled water and once in 500μl 20% (v/v) trichloroacetic acid (TCA). Following 5 min centrifugation at 4000 rpm at room temperature, the cell pellet was resuspended in 100μl 20% (v/v) TCA, glass beads were added and the mixture was vortexed vigorously for 4 min. The supernatant was collected, and the beads were washed twice with 7% TCA to retrieve the remains. The supernatants from all steps were pooled and placed on ice for 45 min. Next, the samples were centrifuged for 10 min at 13000 rpm (TCA precipitation) at 4°C. The supernatant was discarded. The pallets were washed twice with ice-cold 0.1% TCA and then resuspended in a Laemmli loading buffer. Equal protein amounts of each sample were separated by 12% sodium dodecyl-sulfate polyacrylamide gel electrophoresis and stained with Coomassie. The entire protein lane of each sample was cut into three horizontal gel pieces and processed by in-gel trypsin digestion procedure as described before (51) followed by peptides desalting using C18 StageTip (53)

### Native lysis method

Cell pellets were collected by centrifugation of 20 ml culture of each strain in SD media at OD_600_ =1.5. Pellets were washed twice in 1 ml of 25mM Tris pH 7.4, 10mM MgCl_2_, 10% Glycerol, 1mM ATP (BDL) and 1mM DTT (Buffer A). Next, each pellet was dissolved in 200µl Buffer A and 150μl of glass beads were added. The samples were vortexed vigorously for 1 min and then placed on ice for 1 min. This cycle was repeated 8 times in total. After lysis the supernatants were collected using a gel loading tip and centrifuged at 14000 rpm for 10 min at 4°C. The clear supernatants of each sample were collected.

### Native gel, in-gel activity assay

Native lysates (150 μgr) were run in 4% native PAGE in native running buffer (100 mM Tris Base, 100 mM Boric acid, 1 mM EDTA, 2.5 mM MgCl_2_, 0.5 mM ATP (BDL), 0.5 mM DTT) for 2 h at 130 V. For In-gel activity assay, the gel was incubated in 20 ml Buffer A supplemented with 20 μM Suc-LLVY-AMC (R&D Systems) at 30°C for 10 min and imaged under UV light. To visualize 20S and 20S+Blm 10 bands, the gel incubated in 20 ml Buffer A supplemented with 0.02% of SDS for an additional 10 min at 30°C.

### Excision of gel bands of native gels

Active proteasome bands from 4% Native gel were excised under UV light after incubating them with Suc-LLVY-AMC substrate and collected into clean tubes. For different strains, the blade was changed or cleaned thoroughly with 70% alcohol. Excised gel pieces were incubated with destaining solution at RT on shaker overnight before in-gel trypsin digestion procedure as described before (51). The resulting peptides were desalted using C18 StageTip (53)

### Proteasome activity assay on 96-well plate

The Proteasome activity assay was carried out on a 96-well black plate (in triplicate) using the ClarioSTAR plate reader (BMG). Each well was loaded with 30 µL (50 µg protein) lysate with 70 µL of Buffer A buffer containing 20μM Suc-LLVY-AMC (R&D Systems). Assay parameters: Cycle time = 0.36 sec, No. of cycles = 20, Excitation wavelength = 350 nm

### LC-MS/MS analysis

Samples were analyzed using Q-Exactive HF mass spectrometer (Thermo Fisher) coupled to Easy nLC 1000 (Thermo). The peptides were resolved by reverse-phase chromatography on 0.075 × 180 mm fused silica capillaries (J&W) packed with Reprosil reversed-phase material (Dr. Maisch; GmbH, Germany). Peptides were eluted with a linear 60 or 120 min gradient of 6–30% acetonitrile 0.1% formic acid for active proteasomes profiling and whole proteome analysis respectively. In both cases these gradients were followed by a 15 min gradient of 28–95%, and a 15 min wash at 95% acetonitrile with 0.1% formic acid in water (at flow rates of 0.15-0.2 μL/min). The MS analysis was in positive mode using a range of m/z 300–1800, resolution 60,000 for MS1 and 15,000 for MS2, using, repetitively, full MS scan followed by HCD of the 18 or 20 most dominant ions selected from the first MS scan with an isolation window of 1.3 m/z. The other settings were NCE =27, minimum AGC target=8×10^3^, intensity threshold=1.3×10^5^ and dynamic exclusion of 20 seconds.

### LC-MS/MS data analysis

Data analysis and label-free quantification of the whole proteome and proteasome active complexes were carried out using MaxQuant (version Version 2.1.3.0)(54, 55). Proteasomal proteins N-terminal analysis was done based on searches with MSFragger version 3.7 (56) via FregPipe version 19.1 (https://fragpipe.nesvilab.org/). In all searches the raw files were searched against the Uniprot *Saccharomyces cerevisiae* searches of Jan 2020 (that include 6060 protein sequences). MaxQuant searches were performed using tryptic digestion mode with a minimal peptide length of 7 amino acids. Search criteria included oxidation of methionine and protein N-terminal acetylation as variable modifications. All other parameters were set as the default. Candidates were filtered to obtain FDR of 1% at the peptide and the protein levels. No filter was applied to the number of peptides per protein. For quantification, the match between runs modules of MaxQuant was used, and the label-free quantification (LFQ) normalization method was enabled (55) with default settings using LFQ quantification and match-between-runs options. MSFargger searches for proteasomal protein N-terminal cleavage were done by using semi-tryptic digestion. Search criteria included precursor and fragments tolerance of 20 ppm with oxidation of methionine, and protein N-terminal acetylation set as variable modifications.

### Bioinformatics data analysis and statistics

Data analysis was carried using Perseus version 2.0.7 (57). Perseus were used to generate principal component analysis (PCA) plots, hierarchical clustering and volcano plots. Multi sample ANOVA and 2 samples T-tests were done with Perseus using the default settings (S0=0.1, FDR=0.05) unless specified elsewhere. The calculation of the distribution of active proteasome complexes in each strain was based on the sum of MaxLFQ intensities of all the 20S subunits in each sample. For example for WT distribution calculations, the MaxLFQ intensities of all alpha and beta subunits were summed for each one of the 4 complexes (20S, 26S, 30S and 20S^Blm10^) in each one of the biological repeats (n=3). The relative amount of each complex was calculated by dividing its sum of MaxLFQ intensities by the sum of MaxLFQ intensities obtained for all complexes. PIPs distribution calculations were conducted in a similar way but in this case, the relative amount was based on the MaxLFQ intensity of the PIP itself. For Sem1 and Cuz1 the calculations were done using LFQ intensities and not MaxLFQ intensities.

### Structural analysis

The coordinates of the pdb structures 5cz4 (WT yCP), 5fgi (yCPβ1-T1A-β2-T1A:carfilzomib) and 5cz7 (yCPβ5-T1A-K81R:bortezomib)(47) were superimposed as is (no further forced alignment was done) using ChimeraX version 1.3(58). For clarity, only the 20S core is shown. The remnants of prodomain β1 (residues −16 to 0) and β2 (residues −11 to 0) from structure 5fgi and the remnant of prodomain β5 (residues −4 to 0) from structure 5cz7 were highlighted in cyan (β1), green (β2), yellow (β5). The rest of the 20S core is shown in red.

## Abbreviations

CP: Core particle
LFQ: Label-free quantification
MS: Mass spectrometry
PAGE: Polyacrylamide gel electrophoresis
PCA: Principal component analysis
PIPs: Proteasome interacting proteins
RP: Regulatory particles
WT: wild-type

