## Supplemental Figures S1-S8 for "Activity-Guided Proteomic Profiling of Proteasomes Uncovers a Variety of Active (And Inactive) Proteasome Species"

**List of items:**

Figure S1: Proteomic analysis of whole cell extracts.
Figure S2: Proteasome peptidase activity assay.
Figure S3: Comparison of complexes Δα3.
Figure S4: Comparison of complexes ΔSem1.
Figure S5: Comparison of 26S to Lidess 26S in ΔSem1.
Figure S6: Proteasomes and Fub1 amounts in different strains
Figure S7: Representative MS/MS spectra of matured β1, β2 and β5 signature peptides
Figure S8: Representative MS/MS spectra of immatured β2 and β5 signature peptides
Table S1: Whole cell proteomics - Provided in attached Excel file.
Table S2: Whole cell proteomics - Go enrichment analysis of proteins with significant abundance changes - Provided in attached Excel file.
Table S3: Activity-guided profiling of active proteasome complexes (entire lane) - Provided in attached Excel file.
Table S4: Activity-guided profiling of wt active proteasome complexes - Provided in attached Excel file.
Table S5: Activity-guided profiling of Δα3 active proteasome complexes - Provided in attached Excel file.
Table S6: Activity-guided profiling of ΔSem1 active proteasome complexes - Provided in attached Excel file.
Table S7: Distribution of proteasome complexes based on peptidase activity (Quantified by ImageJ) - Provided in attached Excel file.
Table S8: Activity-guided profiling of all active proteasome complexes (separate complexes)- was used to determine the PIPs distribution across proteasome complexes in each strain - Provided in attached Excel file.
Table S9: Semi-tryptic peptides searches of Activity-guided profiling of all active proteasome complexes (separate complexes) - was used to determine the activation state of the proteolytic subunits across proteasome complexes in each strain - Provided in attached Excel file.

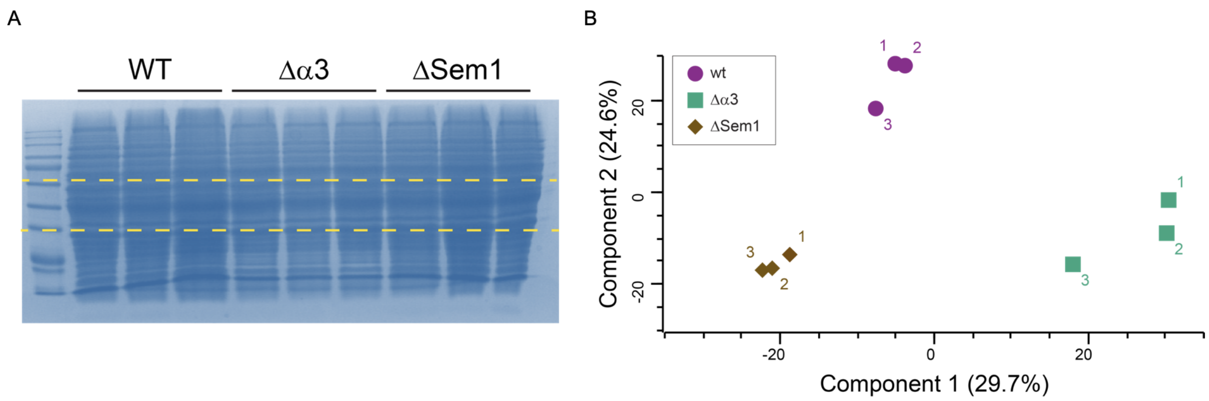


**Figure S1: Proteomic analysis of whole cell extracts. A**. Sample preparation for whole cell proteomics was done following SDS-PAGE separation of whole cell lysate of wt, Δα3 and ΔSem1 cultures. Each lane was cut into 3 regions that were subject to in-gel tryptic digestion**.** **B.** PCA analysis of the LFQ data of identified proteins obtained from the three replicates of each strain.

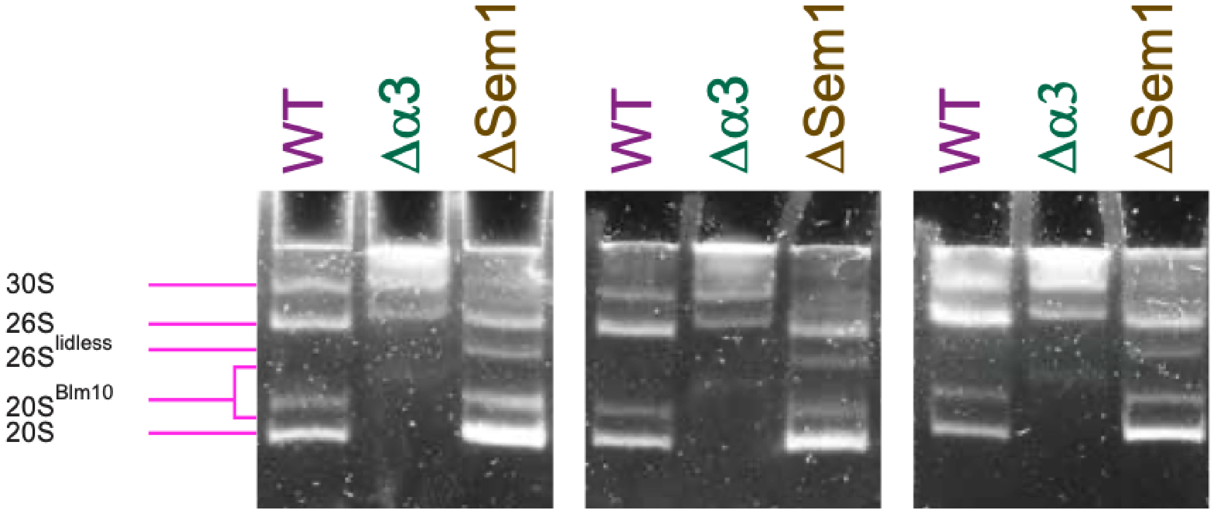


**Figure S2: Proteasome peptidase activity assay** Whole cell native lysate of wt, Δα3 and ΔSem1 cultures were loaded and separated by native-PAGE. Peptidase activity was monitored using the proteasome substrate LLVY-AMC (1).

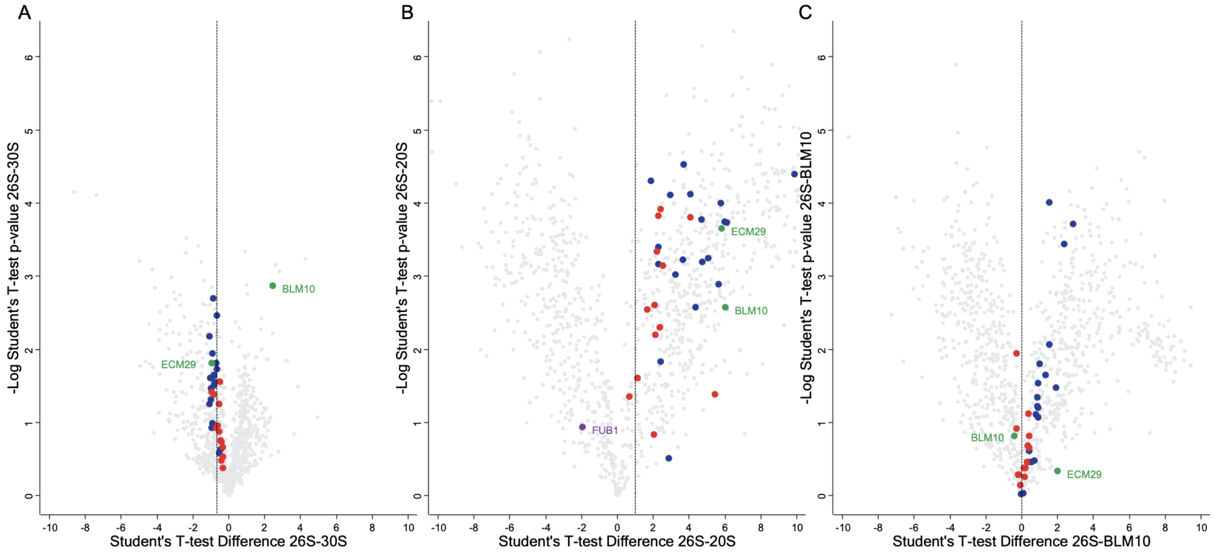


**Figure S3: Comparisons of complexes Δα3 strain. A.** Changes in proteasome subunits and PIPs content of Δα3 26S to 30S complexes. **B.** Changes in proteasome subunits and PIPs content of Δα3 26S to 30S complexes. **C.** Changes in proteasome subunits and PIPs content of Δα3 26S to $26S^{Blm10}$ complexes.

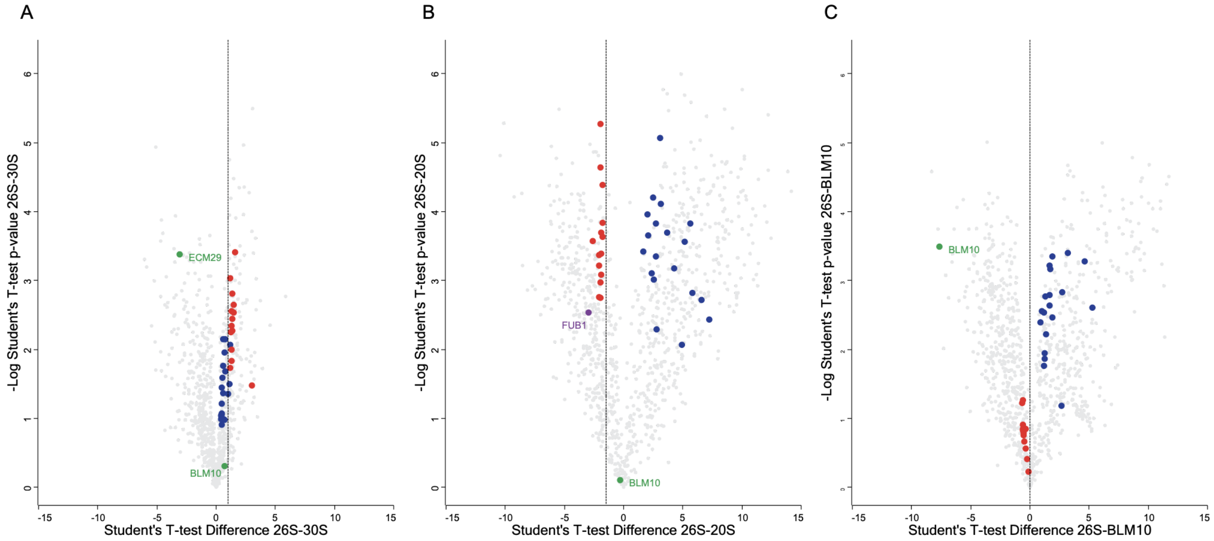


**Figure S4: Comparison of complexes of ΔSem1 strain**. **A.** Changes in proteasome subunits and PIPs content of ΔSem1 26S to 30S complexes. **B.** Changes in proteasome subunits and PIPs content of ΔSem1 26S to 30S complexes. **C.** Changes in proteasome subunits and PIPs content of ΔSem1 26S to $26S^{Blm10}$ complexes.

A

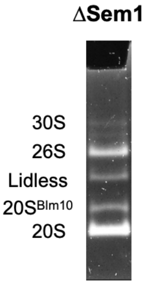


B

| 20S subunit | Fold Change | 19S subunit | Fold Change |
|---|---|---|---|
| α1 | 1.39±0.2 | RPT1 | 1.53±0.09 |
| α2 | 1.45±0.16 | RPT2 | 1.42±0.21 |
| α3 | 1.6±0.42 | RPT3 | 1.21±0.18 |
| α4 | 1.37±0.11 | RPT4 | 1.36±0.06 |
| α5 | 1.35±0.19 | RPT5 | 1.24±0.16 |
| α6 | 1.3±0.31 | RPT6 | 1.11±0.05 |
| α7 | 1.33±0.03 | RPN1 | 1.27±0.14 |
| β1 | 1.28±0.18 | RPN2 | 2.57±0.6 |
| β2 | 1.18±0.07 | RPN5 | 2.04±0.18 |
| β3 | 1.51±0.11 | RPN6 | 2.66±0.51 |
| β4 | 1.57±0.15 | RPN8 | 2.88±0.16 |
| β5 | 1.16±0.21 | RPN9 | 2.47±0.19 |
| β6 | 1.55±0.21 | RPN10 | 3.3±1.37 |
| β7 | 2.01±0.9 | RPN11 | 2.25±0.41 |
| | | RPN13 | 1.26±0.31 |
| | | UBP6 | 1.46±0.17 |
| | | **RPN3** | **7.42±5.24** |
| | | **RPN7** | **23.38±6.45** |
| | | **RPN12** | **23.3±54.48** |

C

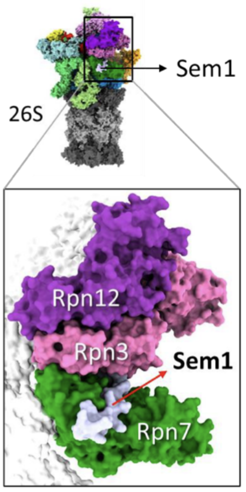


**Figure S5: Comparison of 26S to Lidess 26S in ΔSem1. A.** Native gel shows active proteasome bands from ΔSem1 strain. **B.** The relative fold change of all individual 20S and 19S subunits in 26S proteasome compared to the Lidless-26S proteasome. **C.** Sem1 position in 26S proteasome structure shows direct interactions with Rpn3 and Rpn7 subunits and the proximity to Rpn12. The 26S presentation was generated with ChimeraX(2) based on Yeast proteasome structure PDB code: 6J2X (3).

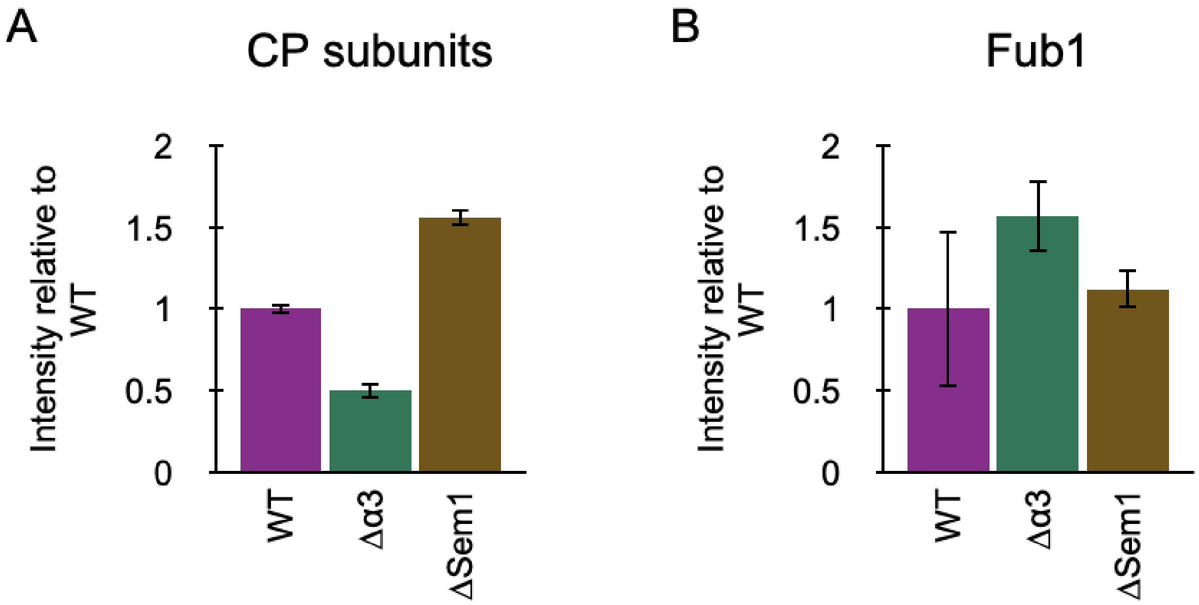


**Figure S6: Proteasomes and Fub1 amounts in different strains . A.** Comparison of the total amounts of proteasomes in the different strains. The LFQ intensities of all CP subunits were summed and the WT total intensity was set as 1 and used as a reference to the comparison to the other strains. **B.** Comparison of Fub1 amount in the different strains. The LFQ intensity of Fub1 in WT was set as 1 and used as a reference to the comparison to the other strains. Note: WT Fub1 LFQ intensities were calculated only in 2 repeats out of 3.

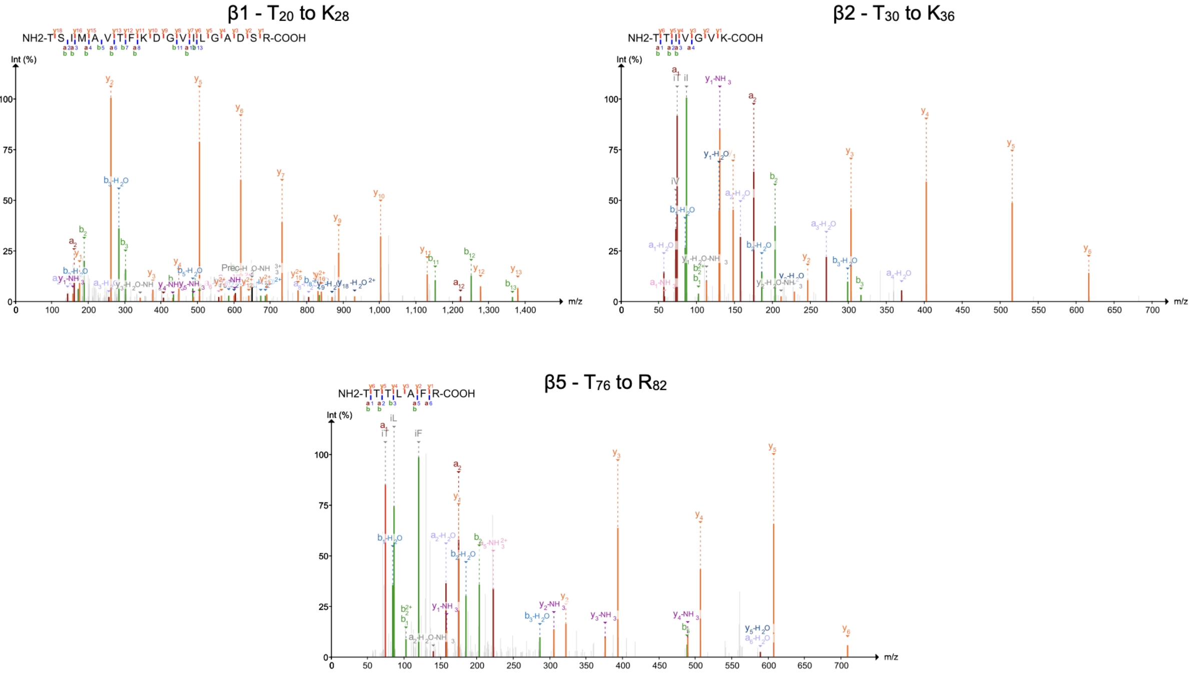


**Figure S7: Representative MS/MS spectra of matured β1, β2 and β5 signature peptides.**

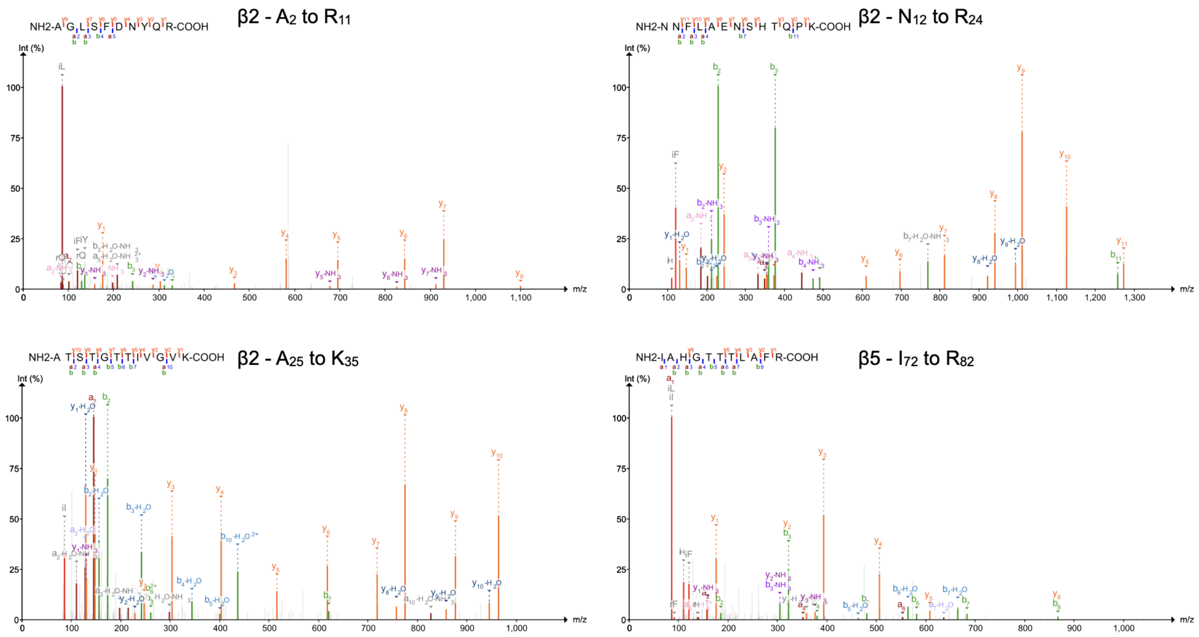


**Figure S8: Representative MS/MS spectra of non-matured β2 and β5 signature peptides.**


<u>References</u>:

1. S. Elsasser, M. Schmidt, D. Finley, Characterization of the proteasome using native gel electrophoresis. *Methods in Enzymology* **398**, 353–363 (2005).
2. E. F. Pettersen, *et al.*, UCSF ChimeraX: Structure visualization for researchers, educators, and developers. *Protein Sci* **30**, 70–82 (2021).
3. Z. Ding, *et al.*, Structural Snapshots of 26S Proteasome Reveal Tetraubiquitin-Induced Conformations. *Mol Cell* **73**, 1150-1161.e6 (2019).
